# Spatially heterogeneous gene flow may hinder linking phylogeographic data to macroevolutionary patterns

**DOI:** 10.1101/2025.11.04.686599

**Authors:** Ethan F. Gyllenhaal, Lukas J. Musher

**Affiliations:** Texas Tech University, Department of Biological Sciences, Lubbock, TX; Cornell University, Department of Computational Biology, Ithaca, NY; Academy of Natural Sciences of Drexel University, Department of Ornithology, Philadelphia, PA

**Keywords:** Biogeography, Coalescent simulations, Gene flow, Microevolution, Macroevolution, Neotropics, phylogenetics, phylogeography

## Abstract

How microevolutionary processes translate to macroevolutionary patterns is a central question in evolutionary biology. Macroevolutionary and biogeographic studies often rely on species trees to make inferences about diversification, but gene flow can lead to incorrect phylogenetic inference. Under a range of biogeographic conditions, gene flow may not occur uniformly across space during diversification, which could lead to macroevolutionary misinferences. For example, whenever a “peripheral” population is relatively isolated and inferred as sister to a “core” clade of populations that exchange migrants, it might solely reflect gene flow, as opposed to macroevolutionary processes such as the order of biogeographic dispersal events. We use simulations of the hypothetical case of spatially heterogeneous gene flow described above and found that relatively low levels of gene flow led to monophyly of the adjacent populations, with longer branches for peripheral taxa, regardless of the true divergence history. We highlight an empirical example wherein lowland antbirds (Family: Thamnophilidae)–a clade known for dynamic gene flow in Amazonia–tend to be young in Amazonia, but older at Amazonia’s periphery. Although it is challenging to know if the simulated bias applies here, our work suggests a distinguishability problem for any nodes stemming from radiations in geographically heterogeneous environments.

## Introduction

How microevolutionary mechanisms driving genetic variation among populations translate to broad-scale macroevolutionary patterns of species richness, age, phylogeny, and distribution is a key question in evolutionary biology [1,2]. Whereas microevolutionary theory posits population divergence and the accruement of reproductive isolation as major processes impacting the accumulation of species [3–5], macroevolutionary theory posits that speciation, extinction, and dispersal drive such patterns [6]. Additionally, microevolutionary theory treats speciation as a prolonged process wherein population divergence by mutation, drift, and selection is greater than any homogenizing impacts of gene flow (i.e., migration) [2,7], while macroevolutionary processes are typically treated as instantaneous. Thus, a key question is, how does gene flow between diverging populations impact phylogenetic inferences when applied to macroevolutionary hypotheses?

One field that aims to bridge micro- and macroevolution is phylogeography, which uses phylogenetic methods to make inferences about how diversification occurs across the landscape [8]. This relies on phylogenies reflecting the true divergence history of lineages within a species or species-complex. For example, if a phylogenetic analysis recovers a group of taxa in one region (e.g., “Region X”) as monophyletic to the exclusion of another taxon in a different region (e.g., “Region Y”), it is commonly assumed that the taxa in “Region X” diverged more recently from each other than the isolated population in “Region Y.” Such patterns can be leveraged to test macroevolutionary hypotheses and examine the interplay between dispersal (range expansion), population subdivision, and speciation [9,10]. However, it is also possible these spatial patterns could be driven by variation in the probability of gene flow across space [11]. Gene flow among diverging lineages has implications for inferring macroevolutionary history and phylogeographic processes in general [12]. Many studies have shown that gene flow between non-sister lineages can confound phylogenetic reconstructions needed for historical biogeographic or phylogeographic analysis [13–17]. Indeed, inferring the evolutionary history of any shared biogeographic pattern requires accurate phylogenetic inferences, which may be biased if gene flow occurs during diversification and is not properly accounted for during data analysis [18,19]. In short, populations that are adjacent in space (henceforth, “core” populations) may be inferred as closest relatives simply due to long-term genetic connectivity, even if more isolated (henceforth, “peripheral”) populations originate from within the “core” (Figure 1). Thus, two questions are: (i) How do spatially heterogeneous rates of gene flow impact phylogeographic inference? And (ii) to what extent might variation in rates of gene flow across space predict broad-scale biogeographic and macroevolutionary patterns shared among co-distributed clades of organisms?

**Figure 1:**
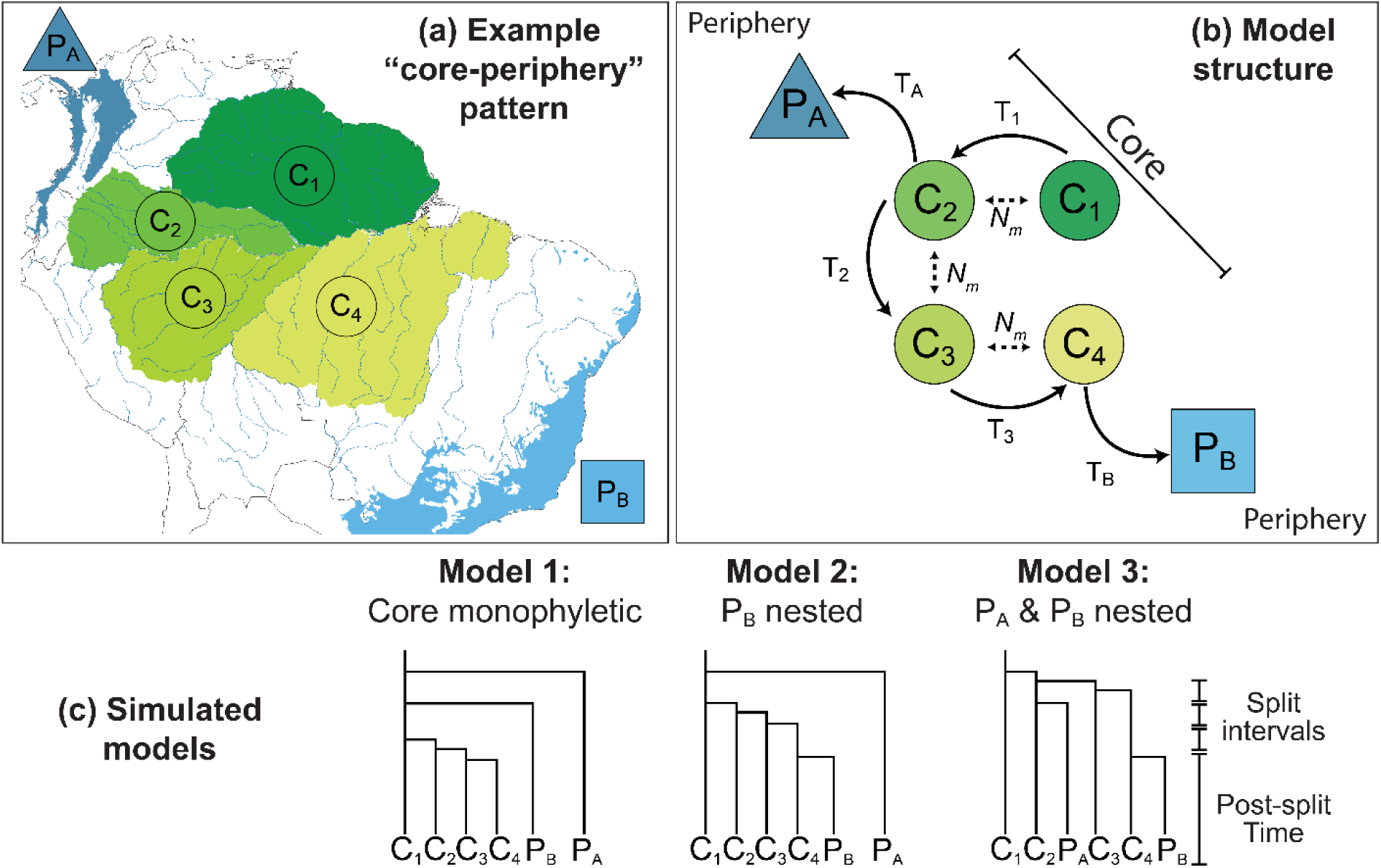
Experimental framework showing (a) the distribution of Neotropical lowland rainforest as an example of populations with higher gene flow among a set of adjacent “core” populations (i.e., Amazonian areas of endemism) compared to two isolated “peripheral” populations (Pacific and Atlantic forests), (b) the structure of simulated models including parameters for migration (N_m_) and split intervals (T), and (c) the three alternative topologies simulated with the model structure in b. Note that this framework reflects a single lineage of allopatric and parapatric (i.e., adjacent and exchanging migrants) populations, which can be interpreted as a single species or complex of three to six species depending on one’s species concept.

Many micro- and macroevolutionary patterns can be produced by spatially heterogeneous gene flow. For example, in rosy-finches (*Leucosticte*), a peripheral clade of populations that occupy islands in the Bering Sea is sister to the North American (i.e., core) continental clade, which experiences high rates of gene flow [20]. A similar pattern is found in Alaskan populations of song sparrows [21]. Although multiple hypotheses including refugial expansions and colonization routes could be true, such patterns can also be viewed as consistent with “peripatric” speciation, a subset of allopatric speciation where peripheral isolates diverge rapidly from a panmictic core range [22,23]. In a more nuanced case, the peripheral island of Makira in the Solomon Islands is likely the most isolated from gene flow due to its relative isolation and unfavorable orientation for receiving migrants [24], but is also the most distant from a common source population, and therefore likely to be colonized last for many taxa [25]. In phylogeographic studies, populations of birds from Makira are recovered as sister to clades of related “core” populations on the other island groups that are adjacent to each other in space and therefore more likely to exchange migrants [26–28]. In these and similar cases, elevated gene flow among some populations and increased isolation of others may mislead biogeographic inference if the impacts of geography on potential gene flow are not considered. This often relies on gene flow between non-sister lineages, but an increasing number of studies have identified this as a potentially common phenomenon, even in cases where secondary sympatry is eventually established [29–32].

Another example that resembles the scenarios outlined above occurs in South American fishes and birds [33–35]. Specifically, some studies have identified young, diversified clades at South America’s “core,” Amazonia, and older (earlier branching) less diversified clades of locally endemic species toward its more peripheral coastal rainforests [35,36]. For example, fish species richness reaches its zenith in the Amazon Basin, which experiences high rates of river dynamics that promote both isolation and secondary contact between fish populations and species. More isolated “peripheral” river basins, however, have reduced species richness, with communities dominated by older, less diversified, but more locally endemic fishes [33]. These patterns in fishes are also similar to those of passerine birds, where Amazonian communities (especially in the highly dynamic western lowlands) tend to have a high richness of young, closely related species, but Pacific and Atlantic rainforests (peripheral) tend to have older lineages and higher regional endemism [36,37]. This core-periphery pattern has primarily been attributed to macroevolutionary processes of instantaneous speciation, dispersal, and extinction. It seems plausible, though, that microevolutionary processes associated with spatial heterogeneity in rates of gene flow might also mimic this macroevolutionary pattern, since the Amazonian landscape is highly dynamic [38,39] and rates of gene flow are partly a function of landscape dynamism [40,41].

At the phylogeographic scale of single species complexes, though, a pattern of monophyletic core taxa and deep-branching peripheral taxa can be challenging to distinguish. For example, some terrestrial groups show monophyly of Amazonian lineages with old divergences from Pacific or Atlantic coastal forest lineages [42–44], others suggest recent colonizations of peripheral forests from within Amazonia [43,45], and yet others show equivocal phylogenetic results (i.e., alternative analytical approaches can yield either pattern; [46]). These disparate results could be influenced by the impacts of gene flow that confound phylogenetic inferences rather than representative of the history of regional colonization and macroevolutionary diversification. This example involves the specific case of South America, but similar patterns of divergence history being clouded by gene flow can be found elsewhere, especially rapid radiations with frequent hybridization such as African cichlids and the Canary Islands’ endemic flora [19,47,48]. Thus it is of key importance to understand how spatial effects on gene flow shape metrics used to assess the drivers of macroevolutionary and biogeographic patterns in conditions where the degree of isolation, and thus probability of gene flow, vary across space.

Here we use simulations at a phylogeographic scale (populations representative of evolutionarily young lineages) to examine how spatial effects on gene flow can shape metrics used to assess the biogeographic and macroevolutionary patterns: monophyletic “core” populations and longer branch lengths for “peripheral” populations. Specifically, we simulate a scenario where some diverging lineages in close geographic proximity (core) exchange migrants with each other but not more geographically distant (peripheral) lineages. We draw inspiration from and model our simulations after South American biogeographic patterns and adopt the term “core-periphery pattern” [35] to describe any biogeographic pattern where some number of [core] species are less isolated from each other in space and recovered as monophyletic to the exclusion of a more isolated [peripheral] species. Specifically, we ask how gene flow among core lineages can mislead phylogeographic inference and result in a monophyletic core with relatively short branch lengths even when peripheral lineages originated from within that core. Although we model our simulations on South American lowland rainforests for ease of conceptualization, these should be viewed as representative of a range of biogeographic scenarios wherein populations that are adjacent in space are more likely to exchange migrants than those more isolated in space, regardless of the phylogenetic relationships among all populations.

## Methods

### Simulation Design

We used coalescent simulations to simulate models that mimic scenarios where gene flow is higher among core populations and reduced for peripheral populations. This framework allowed us to examine how spatially heterogeneous rates of gene flow could impact phylogeographic inferences. Coalescent simulations were performed in msprime v1.0.1 (Baumdicker et al. 2022) and were designed to reflect a scenario where a geographically structured core metapopulation that exchanged alleles served as the source for two isolated peripheral populations (Figures 1A & 1B).

The genomic output of the simulation was a single simulated chromosome with a length of 10 Mbp and recombination rate of 10^-7^ per base pair per generation. The recombination rate was relatively high compared to other birds (e.g., [16,49–52]) to allow for the use of a shorter sequence length for analyses. After each simulation, 2 samples of individuals were output as 4 haploid chromosomes per population as a FASTA file, with one entry per chromosome, with nucleotides modeled using a Jukes-Cantor model and a mutation rate of 4.6 x 10^-9^ mutations/generation (2.3 x 10^-9^ at an arbitrary 2 years per generation) [53].

The demography of the simulation followed a trunk model, where population splits occurred via “budding,” without an impact on the source population (Figure 1C). The four core populations (labeled C_1_-C_4_) exchanged migrants in a linear stepping-stone pattern (i.e., only adjacent populations exchanged genes) at a variable migration rate in terms of migrants per generation (0, 0.2, 0.4, or 0.8). The core populations diverged from population 1 in numeric order [newick=(C_1_,(C_2_,(C_3_,(C_4_)))))]. The two peripheral populations (P_A_ and P_B_) did not exchange migrants with any other populations. For ease of conceptualization and comparison to real-world examples, this model was loosely based on the biogeography of lowland (non-montane) rainforest species in South America, where we allowed gene flow between some, but not all core populations (e.g., the Amazon River mouth is a strong barrier to gene flow for many organisms). In this model, non-Amazonian Pacific and Atlantic coast rainforests are treated as peripheral (Figures 1A and 1B).

We tested three different divergence scenarios for these demographies, allowing peripheral populations to either be nested (arising from within the core) or non-nested (arising from population C_1_ prior to core populations C_2_–C_4_). The scenarios are depicted in Figure 1C and can be described as follows: (1) “ Core nested”– core populations diverged after peripheral populations, (2) “P_B_ Nested”– core populations diverged after P_A_ diverged but P_B_ diverges from C_4_, and (3) “P_A_ and P_B_ Nested”– core populations diverged prior to P_A_, which diverged from C_2_ and P_B_, which diverged from C_4_. For each scenario, population divergences occurred at set intervals, with the core populations diverging sequentially in 3 evenly spaced steps within an interval. In addition to a varying migration rate among simulations, we varied three other parameters: two per-population effective population sizes (50,000 and 100,000), two lengths of post-divergence evolution (200,000 and 400,000 generations), and three lengths of intervals between divergence events (50,000, 100,000, and 200,000 generations). For each combination of parameters, we ran 20 replicates, totaling 3,600 simulations.

### Phylogenetic analyses on simulated data

To examine how spatially heterogeneous rates of gene flow impact phylogeographic inference, we then reconstructed phylogenies for each simulation and assessed the inferred topology relative to the simulated history. Specifically, the output FASTA was used to infer a phylogeny in IQTREE v2.0.3 (Minh et al. 2020) with a Jukes-Cantor model of sequence evolution. We opted to use this method because of its speed, widespread use in phylogenetic analysis, and applicability to the single genomic region simulated, but recognize concatenated methods can perform worse in the face of gene flow [54]. Note that “samples” in this case are single chromosomes (with two individuals and four samples per population).

Output phylogenies were automatically processed using ETE3 v3.1.3 (Huerta-Cepas et al. 2016) in a custom Python script that: 1) removed all but a single outgroup sample, which was then used for rooting; 2) checked if all four samples from P_A_ and P_B_ each were reciprocally monophyletic, labeling the tree “non-monophyletic” if either was not, and subsetting to one individual per population (including each core population) if both peripheral populations were monophyletic; 3) made a copy of the tree with one individual each for the outgroup and populations P_A_, P_B_, and C_1_ and marked the tree as poorly resolved if any bootstrap value in this tree was less than 70 to assess the backbone support; 4) returned to the tree from step 2 and checked if all core samples were monophyletic or, if not, which peripheral populations were nested within it, and then labeling trees accordingly. For trees marked as poorly resolved (i.e., bootstrap < 70) with all monophyletic core samples, we further subsetted the data to one representative per core population and the outgroup, then re-evaluated support values and categorized them as having a monophyletic core. This was specifically to avoid categorizing trees with a polytomy at the base but a well-supported monophyletic core as unresolved. Note that this protocol was not used to parse topologies within those categories. The labels for the trees were then visualized as a heatmap for each parameter combination in R v4.4.1 (R Development Core Team 2023).

To understand how gene flow among adjacent populations might impact relative species’ ages within a clade, we then assessed branch lengths across each simulation’s phylogeny to see how they differed among core and peripheral populations. To do so, we used ETE3 v3.1.3 (Huerta-Cepas et al. 2016) to calculate a variety of branch length statistics using the phylogeny from the end of step 2 above as input. We calculated the distance from each population’s sample to the next split in the tree to get a metric to compare to empirical species “ages” (in substitutions per site), using the mean of the four core populations to represent core divergences. We compiled the means of this metric per parameter combination for use in plotting, as opposed to choosing a single parameter combination.

### Empirical comparison to Neotropical Antbirds

We sampled a species-rich group of Neotropical birds, the Antbirds (Thamnophilidae) to compare our simulated results to real-world data. We chose to use antbirds as an empirical example because they are largely a lowland rainforest radiation, thus allowing direct conceptual comparison to our simulated adjacent “core” and isolated “peripheral” regions. Specifically, there is a high diversity of geographically adjacent species within Amazonia, and reduced diversity in the more isolated humid forests of South America (Atlantic Forest and Pacific Coast Forest). This is in contrast to other groups of suboscines, many of which have exceptional radiations in the Andes, another potential “core” [34,55].We evaluated divergence times for antbird species in three regions corresponding to our simulated core and periphery populations in Figure 1A. We obtained divergence times from a previously published tree [55,56], which we pruned a previously published phylogenetic tree of all suboscines based on the Howard and Moore taxonomy [55,56] to remove all non-antbird tips using the package “ape” v5.0 [57] in R v4.3.1 [58], resulting in a tree containing 236 species. Next we scored antbird geography under four criteria corresponding to Figure 1A. The areas were (1) Pacific coast rainforests and Central America (P_A_), (2) Atlantic coast rainforests (P_B_), (3) regional Amazonia, and (4) widespread Amazonia. Regional Amazonia was scored for species occurring in just one of the following areas: Guiana Shield (population C_1_, Figure 1A), Western Sedimentary Basin (lumped C_2_ and C_3_), or Brazilian Shield (C_4_). Widespread species were defined as occurring in more than one of the three Amazonian regions. Because Amazonian species richness is likely to be highly underestimated [59–61], we treated widespread species as a separate category to help mitigate potential impacts of unrecognized species that are good candidates for future taxonomic splits at smaller spatial scales. We recognize that these discrete categories are somewhat arbitrary, but believe in this case are sufficient to describe the observed pattern we focus on here, regions with different levels of isolation within them containing clear geographic barriers and depauperate antbird communities in the peripheral regions. We then used custom scripts in R v4.3.1 to extract branch lengths from the antbird tree and plot them by geographic category. Finally, we used a Kruskal-Wallis rank sum test with a post-hoc Dunn test and Benjamini-Hochberg correction to evaluate differences in mean species age between Amazonian and peripheral regions using the “dunn.test” function in the R-package FSA [62].

## Results

### Gene flow confounds phylogeographic history and results in monophyly of adjacent populations

Our simulations suggested that relatively low levels of gene flow can produce monophyly of adjacent “core” populations regardless of underlying divergence history (Figure 2). First, simulations wherein the core populations diverged after peripherals (Core monophyletic model) always recovered a monophyletic set of core populations when these topologies had strong statistical support (BS>70 for all nodes). Second, simulations under the “P_B_ nested” model consistently recovered the correct (simulated peripheral nested models) topology when N_m_=0 but tended to recover topologies with a monophyletic core when gene flow was included in the model. This held true for values as low as one migrant every ten generations (N_m_=0.1) when post-split divergence periods were long (T=400k generations). The final set of simulations under the “P_A_ and P_B_ nested” model resulted in fewer well-supported topologies without gene flow and a monophyletic core with gene flow (often with a polytomy of long peripheral branches at the base of the ingroup; Figure S1). We also recovered deeper phylogenetic splits for peripheral than core lineages when gene flow was included in the model, even when peripheral populations arose from within the core. This suggests that the relatively long branches of isolated populations compared to adjacent populations can arise due to elevated gene flow among those populations, rather than in-situ radiation exclusive to a region with geographically structured populations that exchange genes or older populations occupying peripheral regions (Figures 3A, S2).

**Figure 2:**
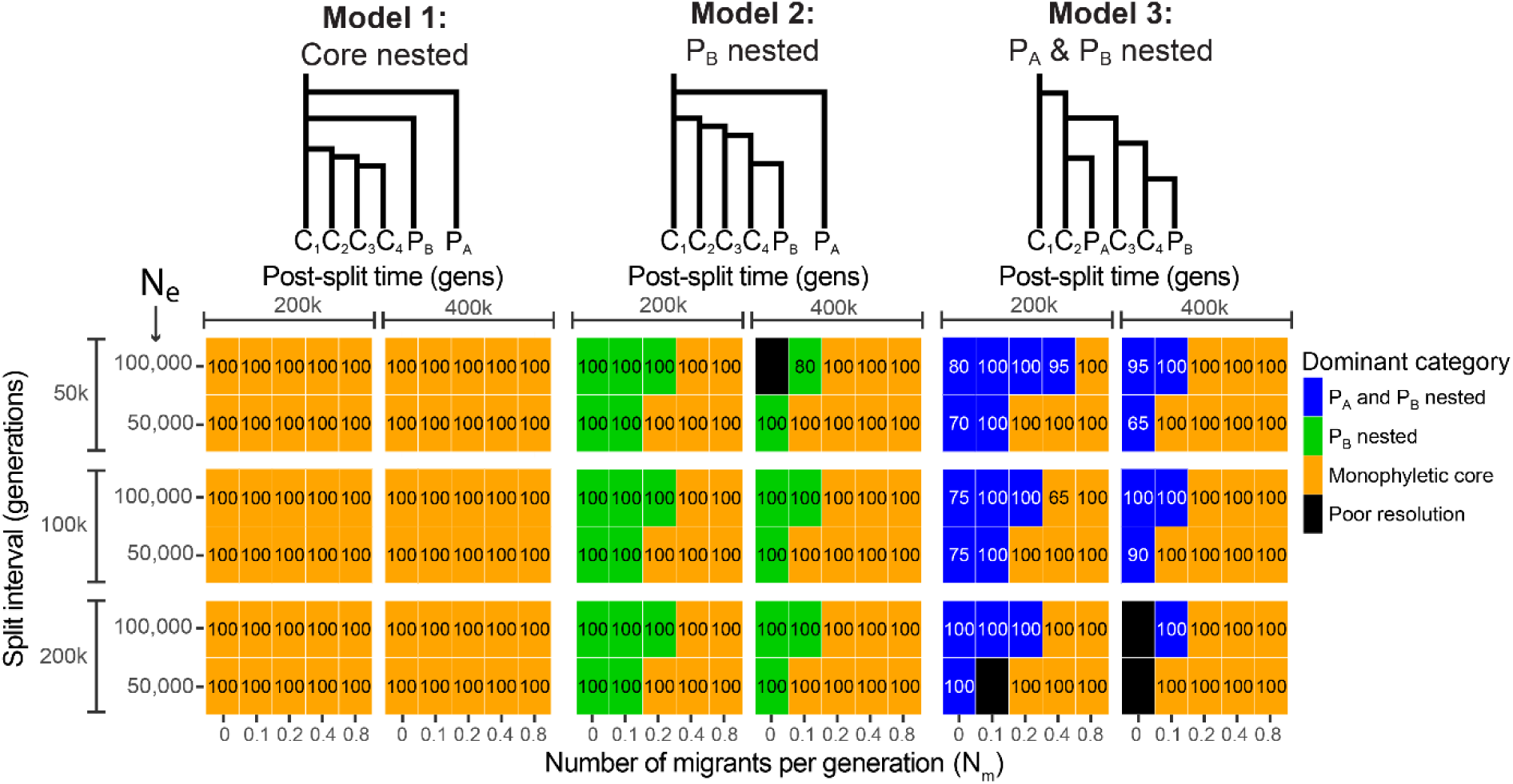
Impacts on phylogeographic inferences. Tile plot summarizing topological results of simulations. For each block of tiles, the x-axis is the number of migrants per generation between adjacent core populations (N_m_; Figure 1), and the y-axis is the effective population size. Among blocks, the x-axis represents the post-split time and the y-axis represents the split interval (see Figure 1). Each tile’s color corresponds to the dominant inferred topology in parameter combinations with mostly well-resolved topologies (>60%). Each tile’s number represents the percentage of all simulations with that dominant topology (not necessarily the simulated topology). All tiles except one (middle column, value of 80) only had one category of resolved topology, so values less than 100 largely represent poorly resolved topologies.

**Figure 3:**
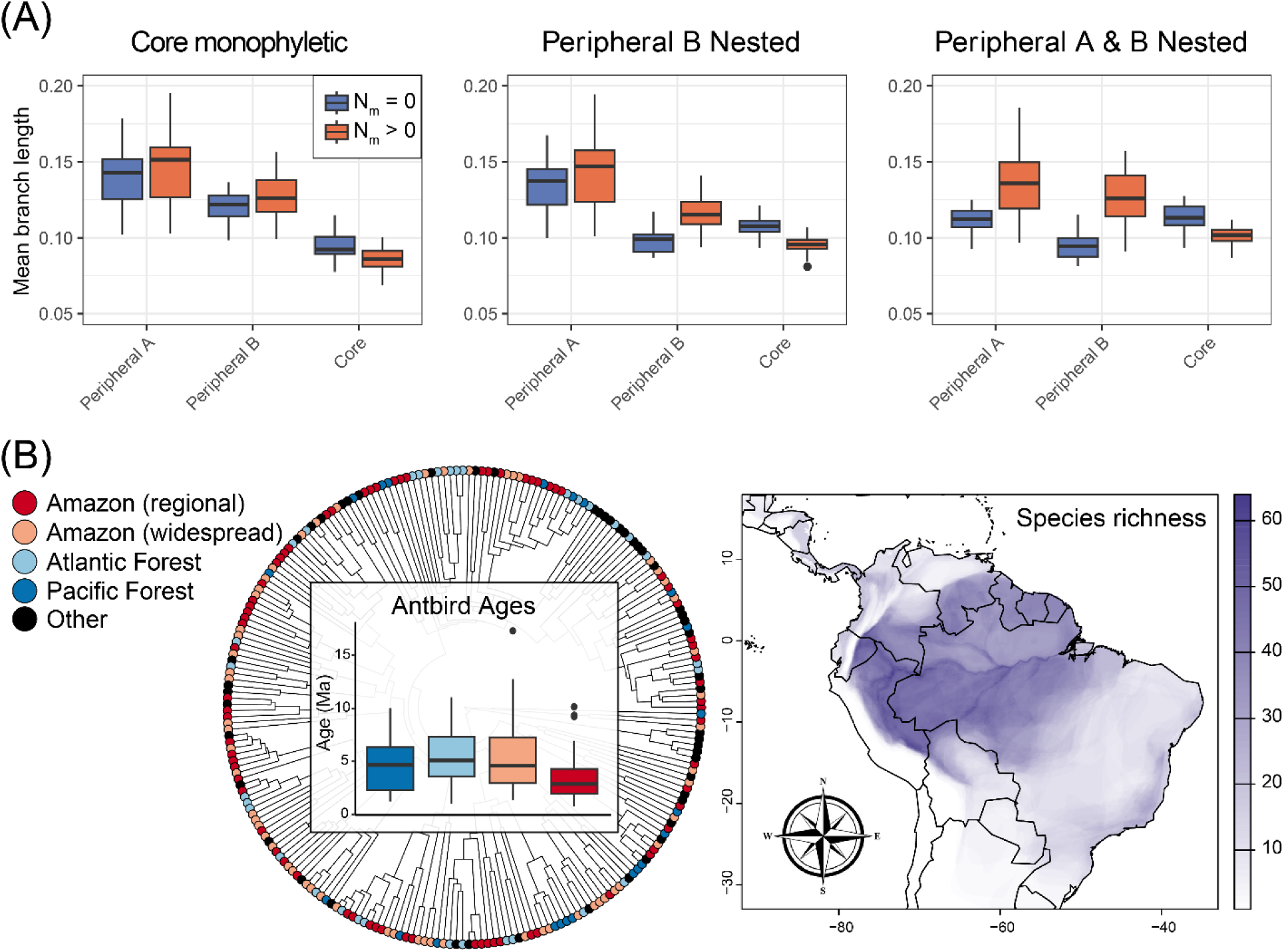
Impacts on phylogenetic branch lengths. (A) Mean branch lengths for three sets of simulated populations showing results for simulations without (left column, blue) and with (right column, orange) gene flow (i.e., Nm > 0). (B) Core-periphery patterns in antbirds (Thamnophilidae): the left panel depicts the ages of lowland antbirds species occurring within the Pacific coast and Central American Forests (dark blue), Atlantic coast forests (light blue), widespread Amazonia (pink), and regional Amazonia (red). The map on the right depicts antbird species richness across South America at a resolution of 0.1 degrees, with darker purple colors representing higher richness. Distributional data are based on BirdLife International species distribution maps (BirdLife International and Handbook of the Birds of the World 2024).

### Amazonian antbirds are younger than their “peripheral” counterparts

Our empirical example of macroevolutionary patterns in Antbirds mirrored our simulations, where geographically adjacent species were younger and more diverse than isolated species at the periphery. Antbirds reach their highest diversity in the western Amazon, a region of high biogeographic-barrier dynamism (Figure 3B)[37]. Species richness declines moving toward the peripheral Pacific and Atlantic coast forests. A Kruskal-Wallis test confirmed that mean antbird divergence times were not equivalent among areas (*X^2^* = 24.2137, *df* = 3, *P* > 0.0001) and a post-hoc Dunn test showed the regional Amazonian species were significantly younger than those in other areas (*P* ≥ 0.02 for all comparisons), but that other areas did not significantly differ (*P* > 0.19 for all). These results qualitatively match our phylogeographic expectations of lineage age derived from our simulations.

## Discussion

We used coalescent simulations to evaluate the impact of spatially heterogeneous rates of gene flow on phylogeographic inference and found that even low levels of gene flow among a subset of populations can severely bias phylogeographic results and interpretations. We used a model based on the assumption that “core” lineages that are adjacent in space should experience more gene flow than those at the periphery either due to closer spatial proximity or increased landscape dynamism, both of which are likely to play a role during the diversification of closely related lineages across the globe. For example, in Amazonia many young bird and mammal lineages are known to be isolated by riverine barriers but experience gene flow across those rivers due to headwater effects, river network dynamics, or insufficient genetic isolation [16,46,63–67]. However, one may expect reduced rates of gene flow between Amazonia (core) and coastal rainforests (periphery), which would require lineages to disperse across the Andes or arid habitats of eastern South America and should be less frequent, especially across the Andes. Though our assumption that peripheral regions experience reduced gene flow relative to core populations seems likely, we suggest that future studies evaluate these expectations more thoroughly using spatially explicit simulations and empirical data. Regardless, our results illuminate the subtle ways in which inferred phylogeographic and macroevolutionary patterns might be influenced by spatially heterogeneous rates of gene flow between non-sister lineages. Moreover, these results agree with classic population genetic theory, that relatively few migrants per generation can lead to substantial genetic homogenization [68]. As antbirds exemplify a pattern of younger lineages that are adjacent in space (Amazonian) and older lineages that are more isolated in space, a pattern also recovered for many types of organisms [33,36], future studies should test whether this pattern might be explained, in part, by increased gene flow among non-sister Amazonian species occurring in close geographic proximity [40].

### Implications of spatially heterogeneous gene flow for phylogeographic inference

Gene flow impacts evolution in multiple ways, including as a creative force that leads to adaptive change or as a destructive force that reduces divergence between populations [69–72]. For example, during diversification, gene flow that occurs between non-sister taxa can deplete the historical phylogenetic signal from the genome by introducing new gene histories that differ from the historical branching pattern [14,16]. This process of diversification with gene flow leads to heterogeneous signals in the genomes of diverging populations, which complicates phylogenetic analysis, and thus any downstream inferences that hinge on phylogenetic relationships. This effect is known [14,17,51], but here we demonstrate the additional challenge posed by the biogeographic context of diverging lineages wherein spatially heterogeneous rates of gene flow can produce biased results of interest for testing biogeographic hypotheses. This is a problem if interpreted at face value, the common topology across taxa could be seen as support for a single biogeographic hypothesis regardless of the true divergence history.

The role of spatially heterogeneous rates of gene flow in shaping phylogenetic inference should be carefully considered by biogeographers, particularly those working on recently diverged taxa. Gene flow is largely a function of the geographic contact between diverging lineages, so rates of gene flow among populations should be highest in regions where many young lineages occur in close proximity, including where biogeographic barriers are most dynamic. In this way, a dynamic landscape simultaneously impacts rates of lineage formation, dispersal, and extinction [34], but also rates of gene flow among these young diverging lineages [64]. Moreover, because core lineages tend to be younger (e.g., Albert et al. 2017; Crouch et al. 2019; see also Figures 3 and S2), they may be more likely to experience gene flow because young lineages are expected to lack intrinsic reproductive isolation [66], though this was not examined here. Low levels of gene flow can compound these effects by hindering the opportunity for reproductive isolation to arise [5,73] and the analytical effect of shortening inferred branch lengths and therefore species ages [14,17,51,74] (Figure 3). Although in a neutral model, proximity is primarily an opportunity for gene flow, in natural populations it can also be an opportunity to complete speciation via reinforcement [4,75,76] or character displacement [77,78]. Thus, gene flow among adjacent core populations might inhibit diversification in some cases, but could also facilitate ecological divergence, reproductive isolation, and eventually sympatry in other cases, thereby increasing local diversity over time [5]. Indeed, for the levels of migration we simulated, each individual core subpopulation was typically recovered as reciprocally monophyletic, demonstrating that population divergence can be maintained under the same migration regimes that can bias deeper level phylogenetic inference.

We model our simulations in part on core-periphery macroevolutionary patterns in South America, but the process we simulate is applicable to a broad range of studies interpreting recent-time phylogenetic data. For example, gene flow might impact the interpretation of comparative studies, where a consistent phylogenetic pattern is driven by relative microevolutionary isolation, not shared biogeographic origins and divergence histories alone.

This is because the same processes that drive microevolutionary histories of isolation and gene flow, also drive speciation and dispersal, respectively. Thus, the same macroevolutionary patterns can emerge despite different underlying microevolutionary processes. Simply put, whenever a population is relatively isolated from others and inferred as sister to a clade of populations that do exchange migrants, it may be reflective of microevolutionary processes such as gene flow and drift, as opposed to macroevolutionary processes such as the order of biogeographic dispersal events or differential diversification rates. These implications are wide-reaching as it is safe to assume that gene flow always occurs heterogeneously across space as geographic barriers to gene flow always vary in size, extent, and dynamism [41,79]. We note that although gene flow is not necessarily driving all examples wherein adjacent populations are monophyletic to the exclusion of peripheral populations, it is important to consider whether or not alternative scenarios are distinguishable in the face of low levels of gene flow within a given dataset.

We simulated relatively long sequences with high recombination, allowing migrant alleles to become incorporated into the sequence stochastically over generations. This gene flow predictably altered the inferred phylogenetic topology (Figures 2 & S1). Although many phylogeographic studies test for gene flow, at times explicitly accounting for it using networks or similar analyses [52,80], many common tests, such as those derived from the D-statistic (ABBA-BABA)[81], are typically performed as post-hoc tests assuming a species or population tree to identify potential instances of gene flow among branches in that tree. Thus, tests of gene flow utilizing this incorrect topology could be positively misleading as they would assume the wrong history. Although some alternate methods that incorporate uncertainty such as sampling a posterior distribution of tree topologies for biogeographic analyses may be productive (as in [82]), if the signal is sufficiently strong and data volume high the biased signal may overwhelm the true branching order.

We suggest that future phylogeographic work use additional methods to infer the true branching history of populations before estimating gene flow. For example, analyses that account for genome architecture, including recombination, selection and sex-chromosome evolution, or simultaneously estimate phylogenetic history and introgression would be helpful to appropriately reconstruct phylogeographic history when focal taxa experience gene flow [16,49–52]. However, we caution that complex patterns of past and present introgression–particularly with ghost lineages–may render the “true” branching pattern lost across most of the genome [14,83,84].

Future simulation work might aim to explore what methods are best at both untangling complex histories of gene flow and accurately reporting uncertainty due to this gene flow. For example, it might explore alternate data types, such as using ancestral recombination graphs at the phylogeographic scale, although their utility might be limited to the phylogeographic scale and below [85]. Nevertheless, dissecting competing genome-wide signals of gene flow and divergence, can tell a more detailed biogeographic story, wherein the spatial context of population isolation and connectivity lead to multiple phylogeographic signals across the genome, and thus superimposed histories of lineage fission and fusion [41].

### Might a dynamic Amazonian landscape promote species persistence?

We highlighted an empirical example wherein species ages and diversity of antbirds vary non-randomly across the South American continent. Although we speculate that these patterns might be partly attributable to histories of elevated gene flow in Amazonia relative to peripheral regions, the results are also relevant to understanding the history of the Neotropics in general. A key theme in Neotropical diversification research is that Amazonia accumulated high species richness due to low extinction rates relative to temperate regions [55,86,87]. This has typically been attributed to relative stability of the Amazonian climate, especially in the hyper-diverse western Amazon, which enabled high rates of species persistence. An additional, non-mutually exclusive hypothesis is that low extinction and species persistence are associated with a *dynamic* landscape that allows species to maintain population connectivity and expand distributions over time. In the same manner that large ranges can buffer against extinction [88–90], low levels of dispersal can allow for both re-colonization of the ranges of extinct members of a species-complex and genetic rescue of populations under a changing environment. Under this hypothesis, persistence is a function of spatial factors that, although not necessarily independent of climate and range size, are fundamentally different. Rather, increased persistence of populations may partly be a function of long-term gene flow among Amazonian populations due to landscape dynamism rather than strict climatic stability (e.g., Albert et al. 2017; Musher et al. 2022, 2024). Although we do not explicitly test this hypothesis in the present study, our analysis of data from antbirds is consistent with its expectations, suggesting additional study is needed. For instance, a dynamic Amazonian landscape should result in high levels of young lineages in this region, which we have found for Amazonian antbirds (Figure 3). Gene flow between these young Amazonian lineages might also artificially reduce estimated divergence times, which may contribute to the significant difference in divergence times among core and peripheral biogeographic areas. Moreover, a dynamic landscape also allows populations to expand their ranges over time; we found that widespread Amazonian antbird species tend to be older than regionally distributed species. We suggest that this hypothesis needs further testing at both micro- and macroevolutionary scales for terrestrial Amazonian taxa [41,59]. For example, future simulation work might blend microevolutionary and macroevolutionary simulations to determine the role of gene flow occurring early in divergence could play in shaping inferred phylogenies at deeper time scales.

## Data Availability

The code, input files, and several output files used in this study are available at https://doi.org/10.5061/dryad.9w0vt4bsb. Additionally, a subset of the code is available at https://github.com/ethangyllenhaal/CorePeripheralGeneFlow.

## Author Contributions

EFG and LJM conceptualized the project, conducted the analyses, and wrote the paper.

## Acknowledgements

We thank J.D. Weckstein, E. Griffith, J. Merwin, K. Kuabara, M.A. Pérez-Pérez, J.D. Manthey, J.P. Hruska, and M.C. Tocora for comments that improved the quality and clarity of our manuscript. We would like to thank the UNM Center for Advanced Research Computing, supported in part by the National Science Foundation (NSF), for providing the high-performance computing resources used in this work.

