## Supplementary material for "Spatially heterogeneous gene flow may hinder linking phylogeographic data to macroevolutionary patterns": Figure S1; Figure S2

**Spatially heterogeneous gene flow among core and peripheral lineages impacts phylogeographic inference**

**Figure S1: Core-periphery impacts on phylogeographic inferences and deeper phylogenetic resolution**. Tile plot summarizing topological results of simulations, identical to figure 2 in input data except with different qualifications for poor resolution. In this figure, poor resolution includes cases of poorly resolved backbones, regardless of if the core populations are inferred as monophyletic. For each block of tiles, the x-axis is the number of migrants per generation between adjacent core populations (m; Figure 1) and the y-axis is the effective population size. Among blocks, the x-axis represents the post-split time and the y-axis represents the split interval (see Figure 1). The base color corresponds to the dominant topology in simulations with well-resolved topologies (>60%), and the number represents the percentage of all simulations with that dominant topology (not necessarily the simulated topology). All tiles except one (middle column, value of 80) only had one category of resolved topology, so values less than 100 largely represent poorly resolved topologies.

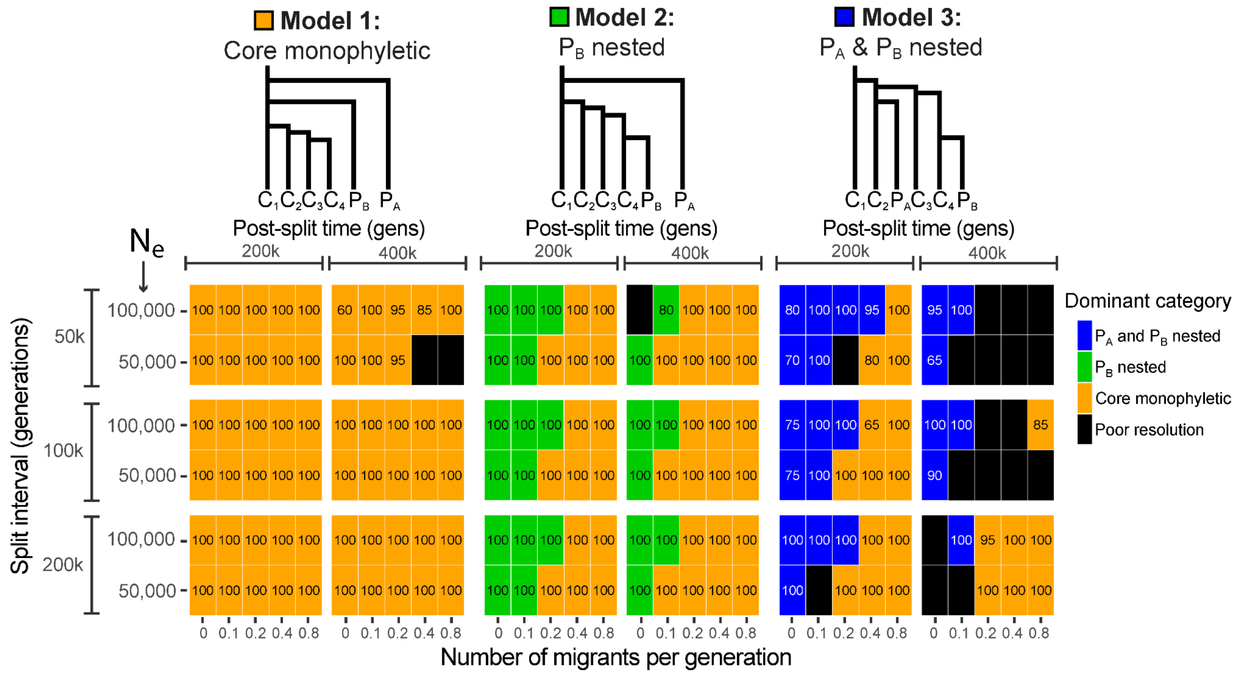

**Figure S2:** Tile plot of the ratio of the mean branch length among core populations to the minimum branch length for either peripheral population as a proxy for population divergence as in Figure 3. The color represents the value of this ratio, with values greater than one (with elevated transparency) largely corresponding to non-monophyly of the core populations, and values less than one corresponding to different levels of divergence among the core populations.

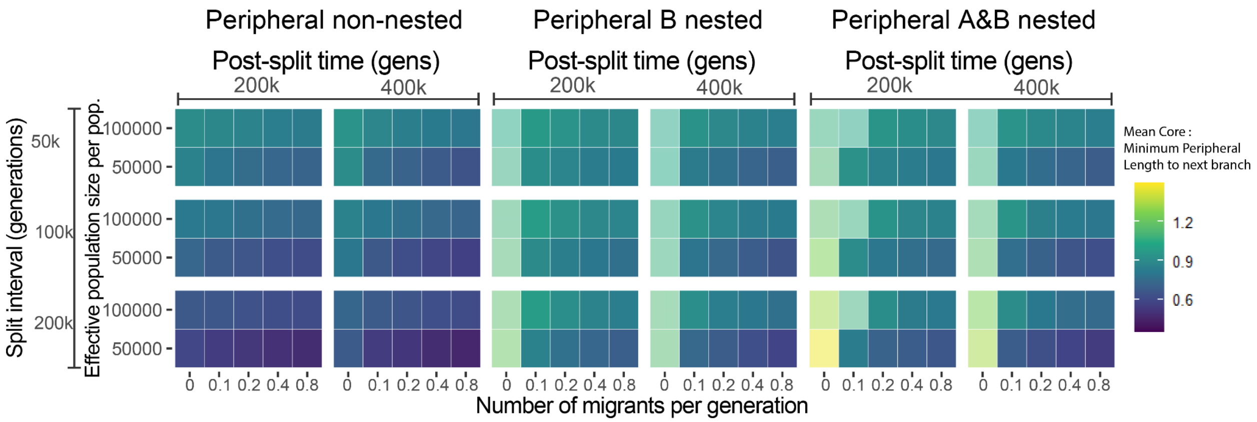
